# Vitamins in native tree pollens are essential to support fitness of wild-caught *Bombus* spp. queens, with positive carry-over effects

**DOI:** 10.64898/2026.08.14.744728

**Authors:** Mathilde L. Tissier, Sarah MacKell

**Affiliations:** Université de Strasbourg, CNRS, IPHC UMR 7178, F-67000 Strasbourg, France; Département Des Sciences Biologiques, Université du Québec à Montréal, Montréal, Québec, Canada; Wildlife Preservation Canada, 5420, Suite 234, Hwy 6, Guelph, ON N1H 6J2, Canada

**Keywords:** nutrition, fitness, wild bees, conservation, vitamins

## Abstract

To implement adequate nutritional landscapes to bumble bees and inform conservation efforts, we need a better understanding of how different pollen sources impact bumble reproductive performances and phenology. We assessed the impact of multiple native pollen on bees’ reproductive success by feeding 40 wild-caught *Bombus impatiens* and *B. griseocollis* queens with either commercially wildflower pollen mix commonly used to breed *Bombus* queens (control group) or 75% red maple pollen (*Acer rubrum*) and 25% pollen mix (RM group). In a second experiment, we monitored the reproductive success of 45 microcolonies of workers, harvested from *B. impatiens* colonies. Microcolonies were fed either: 1) pollen mix (N=15), 2) pollen mix and hawthorn (*Crataegus canadensis*, N=15), or 3) pollen mix and sumac (*Rhus typhina*, N=15). In experiment 1, RM significantly increased reproductive outputs for both species. RM-fed founding queens were twice more likely to produce workers, and produced them significantly earlier, though total number of workers did not vary between groups. However, RM-fed founding queens produced significantly more brood cells, males and gynes than control colonies, and produced them significantly earlier. In experiment 2, we found an interaction between initial colony diet and current microcolony diet. Overall, workers harvested from RM-fed colonies produced significantly more brood cells and males than workers harvested from control colonies. However, when fed hawthorn pollen during experiment 2, workers from control colonies performed as well as workers from RM-fed colonies. This suggests carry-over effects from early pollen sources provided to colonies on workers, which may be circumstantially offset by pollen sources with specific nutrient profile. Using detailed nutritional information of each pollen (including data on 32 nutrients), we provide nutritional requirements of pollen to support bumble bee reproduction, especially regarding vitamin B3 and vitamin C contents. This study will help inform land managers and captive breeding programs on the ideal pollen sources for bumble bees to ensure reproductively successful colonies.

## Introduction

Insect pollinators, including wild bees, are declining worldwide (Cornelisse et al., 2025; Wagner, 2019). Among pollinators, bumble bees face particularly high risks of extinction (Cornelisse 2025). Threats to pollinators are multiple and include habitat loss, climate change, pesticides and disease spillover from managed bees, as well as malnutrition (Cameron & Sadd, 2020; Cornelisse et al., 2025; Guzman et al., 2024). Recent work highlights that those threats to pollinators may have additive and non-linearly interactive effects (Birkenbach et al., 2024; Tissier et al., 2025).

Access to diversified flower resources or nutritious pollen has been shown to improve pollinator response and resilience to environmental stressors like heat waves and pesticides (Birkenbach et al., 2024; Cameron & Sadd, 2020; Costa et al., 2022; Klaus et al., 2021; Tissier et al., 2025). Maintaining a diet of high nutritional quality, balanced in essential nutrients, supporting bees’ resilience to such environmental stressors is thus vital. Increasing evidence supports the idea that, more than food quantity (e.g. quantity of flowers, pollen and nectar available), the nutritional quality of dietary resources is essential to support bee health, social behavior, reproductive performances and survival (Birkenbach et al., 2024; Parreño et al., 2022; Vaudo et al., 2024). This includes the ratio of proteins to lipids (P:L ratio), total content in essential amino acids (EAAs), or content in fatty acids or B-complex vitamins (Jovanovic et al., 2021; Vaudo et al., 2024). However, while extensive information is available on the nutritional requirements of managed European bees like *Apis mellifera* and *Bombus terrestris* (Barraud et al., 2022) and extensive progress has recently been done on Northern American managed bumble bee *B. impatiens* (Vaudo et al., 2016), or a couple of *Osmia* spp solitary bees (Barraud et al., 2022; Yourstone et al., 2021), still little is known on the nutritional requirements of wild bees, including most bumble bee species. Moreover, studies looking at the dietary requirements of bumble bees are often conducted on queenless microcolonies of workers (Barraud et al., 2022; Genissel et al., 2002; McAulay & Forrest, 2019; Moerman et al., 2017; Tasei & Aupinel, 2008), and may not fully represent the nutritional requirements and energy trade-offs of founding queens (Wynants et al., 2022). Such knowledge gaps are limiting our ability to implement effective conservation measures supporting diversified habitats meeting the nutritional requirements of a diversity of bee species (Barraud et al., 2022; Vaudo et al., 2016).

Designing conservation habitats supporting wild bumble bees and meeting their nutritional requirements is receiving considerable effort (Torchio et al., 2024; Vaudo et al., 2016), especially in agroecosystems, where pollinators are rapidly declining, while also being most essentials through their pollination services. Agroecosystems relying on monocultures are often nutritionally deficient, with damaging effects on bumble bee behavior, reproduction and survival, emphasizing the need for targeted actions to restore nutritional balance, beyond mere functional diversity. For instance, organic monoculture of squash (*Cucurbita pepo*) reduced the reproductive performance and colony growth of commercial *B. impatiens* (Gauger et al., 2025). Corn and dandelion, on the other hand, are known to be deficient in niacin (vitamin B3) and its precursor, the essential amino acid tryptophan, leading to aggressiveness, reduced survival and oophagy in the European *B. terrestris* (Genissel et al., 2002; Tissier et al., 2023) and reduced reproductive success in the Asian *B. eximius* (Wang et al., 2025). Identifying plants whose pollen is nutritious to bumble bees and can support reproduction and survival throughout the season but also from year-to-year is therefore of paramount importance to be able to design nutritionally adapted conservation habitats. While nectar is the main source of food for adult bumble bees, the composition of pollen is determinant for colony performance and reproductive output. Early flowering trees such as willows, maples and oaks are key resources providing important nectar and pollen provisions for social and solitary bees (Batra, 1985; Chrzanowska et al., 2024; Yourstone et al., 2021), including in farmland habitats. However, we lack information on whether they support the development of wild bumble bee colonies and whether all *Bombus* spp species show the same response. We previously found that consumption of red maple (*Acer rubrum*) – whose tissues are particularly rich in niacin or vitamin B3 (Burkholder & McVeigh, 1945) – increased reproductive success in a wild vertebrate (Tissier et al., 2020). Based on this finding and on evidence that red maple pollen offers important resources to pollinators early in spring, we designed this study to improve our knowledge on the nutritional value of flower pollen native to North America on the reproductive performances of wild native bumble bees. We first investigated the benefits red maple pollen on the longevity and reproductive output of 40 wild caught overwintered bumble bee queens of two species, *B. impatiens* and *B. griseocollis*. We then investigated the nutritional properties of hawthorn (*Crataegus canadensis*) and sumac (*Rhus typhina*) on the reproductive output of queenless microcolonies of *B. impatiens* workers from the previously caught wild founding queens.

Red maple pollen is nutritionally rich (Chrzanowska et al., 2024; Tissier et al., 2023). We thus expected that queens fed red maple pollen would exhibit improved reproductive outputs (i.e. increased probability to initiate a colony, greater colony size and strength, increased production of reproductive and that adults would emerge earlier) and have greater longevity than control queens fed a wildflower pollen mix. Similarly, we predicted that hawthorn and sumac would be more nutritious than wildflower pollen mix, thereby improving performances (reproduction and survival) of microcolonies fed with these pollens.

## Material and methods

### Queen collection and rearing

Between May and early June 2021, we collected 24 *B. impatiens* and 20 *B. griseocollis* queens at 19 different sites across Southern Ontario (Figure S1). Upon collection, bumble bee queens were placed in individual, small, and well-ventilated vials (Rowe et al., 2023), provided with nectar and put in a cooler to maintain them at low temperature and reduce activity until installation in the lab (Rowe et al., 2023). Installation was done at the Bumble Bee Conservation Lab of Wildlife Preservation Canada located at African Lion Safari (Ontario, Canada) within 24 hours of capture and left undisturbed for 48-72 hours. Installation was conducted according to protocols used by the US Department of Agriculture at the Pollinating Insect—Biology, Management, Systematics Research department in Logan, Utah, who have successfully reared wild-caught bumble bee queens for several years, including rare and declining species such as *B. occidentalis* (Rowe et al., 2023).

Wild-caught queens were installed in individual initiation boxes (15 x 15 x 10 cm, BioBest, Canada, following a detailed published protocol (Rowe et al., 2023)) and transferred to a larger colony box (29 x 22 x 13 cm, BioBest, Canada) once they produced 10 or more workers. Upon installation, each queen was provided with ad-libitum nectar substitute and pollen. The nectar substitute consisted of a 50% cane syrup solution with preservatives, prepared as previously described (Rowe et al., 2023). Pollen was provided in the form of pollen balls, prepared as described in (Rowe et al., 2023): pollen loads were grounded and mixed with nectar substitute to make a dough, that was then cut into that was then cut into uniform 0.40 ± 0.10 g portions. Queens that had not produced any offspring yet were given pollen balls that were covered in previously heated beeswax, whereas those with offspring were given unwaxed pollen balls (Rowe et al., 2023). New pollen balls were added every other day, while nectar was changed every 4 days. Colonies were maintained under red lighting at 26 ± 2°C and 55-65% humidity, upon installation and throughout the season and experiments.

Although survival and installation success were high overall, four *B. impatiens* queens died within the first week after being installed, leading to a final sample size of 20 queens for each species. These four queens were not included in the experiment.

### Experiment 1 – pollen effects on performances of wild-caught bumble bee queens

To investigate the effects of early-spring pollen on reproductive and survival performances, we conducted a first experiment on 40 wild-caught overwintered bumble bee queens (20 *B. impatiens* and 20 *B. griseocollis*). Upon installation, all queens were fed 50% sucrose solution and commercial mix of wildflower pollen (commercial mix no 17222, wildflower pollen, *Plant Products*, Canada) for a week, after which each queen was randomly attributed to an experimental diet while considering their date of capture to avoid having all early queens on the same diet (day 1 of the experiment). Half of the queens of each species were then fed a red maple pollen diet (N=20 queens, 10 per species; “Red Maple Mix”, hereafter RM), while the other half was maintained on the commercial mix pollen diet (N=20 queens, 10 per species; “Commercial Mix”, hereafter CM).

Pollen balls for the RM diet were prepared as detailed above and were composed of 75% monofloral red maple pollen (RM; 100% red maple) and 25% commercial pollen mix (CM1; composed mostly of clover and dandelion). This 75-25% ratio was implemented to investigate the nutritional value of red maple pollen on queen performances, while 1) seeking to mimic natural conditions in which bees are unlikely going to feed several weeks on a single pollen source and 2) preventing the deleterious effects of a 100% monofloral pollen diets compared to multi-species mixes (McAulay & Forrest, 2019; Moerman et al., 2017). To limit the risk of a nutritional stress while changing the diet, RM-fed queens were provided with a 50-50% mix of red maple and commercial mix during 7 days before switching to the 75-25% maple-mix pollen (Figure 1).

**Figure 1:**
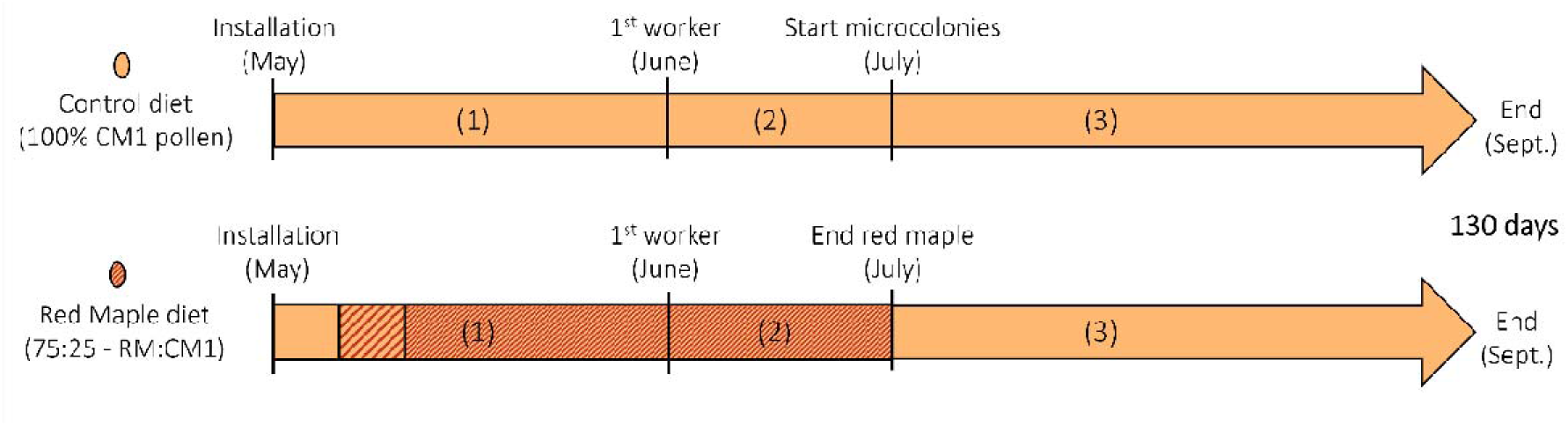
timeline of experiment 1 with feeding patterns for both the control and red maple diet groups. In period (1) colonies were provided with one waxed pollen ball (0.4g) from their respective diets every other day; then (2) they received one additional unwaxed pollen ball from their respective diets for every additional two workers. Finally (3) they all received unwaxed wildflower mix pollen (100% CM1). The control group thus received wildflower mix throughout the 130 days of the experiment. The Red Maple group received wildflower mix upon installation (CM1 pollen), followed by a mix of red maple (RM pollen) and wildflower mix (with a 50:50 ratio first (yellow with red stripes) and then a 75:25 ratio (yellow with narrow red stripes), and was then fed 100% wildflower mix until the end of the experiment. We started harvesting workers in July to initiate microcolonies for experiment 2 (*Start microcolonies* on the figure).

Each queen was respectively provided one CM1 or RM pollen ball of 0.4 ± 0.10 g on day 1 of the experiment. Pollen balls were then changed every other day, except if the queen started to lay eggs on it or if it was attached to something in the box. Upon emergence of the first worker, each colony was provided with an additional pollen ball, of the commercial mix for the CM1 group and of the 75-25% maple-mix for the RM group. This quantity was then gradually increased over the experiment, with one additional ball of commercial mix provided every other day per two new workers emerging in colonies of all diet groups (phase 2, Figure 1). Once the first worker emerged, bees were fed with unwaxed pollen dough (prepared as detailed above, minus the wax) and were supplied new pollen based on consumption to ensure ad-libitum access to pollen. We stopped providing red maple pollen to all queens from the RM diet group in July to mimic natural cycles of pollen availability. They were subsequently fed wildflower mix (Figure 1). As not all queens were installed into the lab at the same time, this made the RM pollen group receive the maple-mix pollen for 7±1 weeks: the 20 colonies of the Red Maple pollen group received the maple-mix pollen for 45±3 days (*B. griseocollis*) and 59±1 days (*B. impatiens*). This difference is reflective of *B. griseocollis* later emergence date compared to *B. impatiens*. Indeed, *Acer* trees flower early in the season, and by emerging on average 2-3 weeks later than *B. impatiens*, *B. griseocollis* founding queens are exposed for shorter duration to *Acer* pollen and are less likely to store large amounts of red maple pollen compared to *B. impatiens*. RM colonies that needed over three balls of 0.4g of pollen were given commercial mix pollen on top of three red maple pollen balls to ensure pollen quantity was not a limiting factor. The RM colonies were switched to the commercial pollen mix starting July 18^th^, until the end of the experiment on September 28^th^.

Every other day, from queen installation to the end of the experiment, we monitored the following parameters to assess performance on diet: (i) number of brood cells, (ii) number of workers, (iii) number of males, (iv) number of gynes and (v) founding queen longevity. These data enable both an assessment of overall queen/colony performance on diet during the experiment, and an evaluation of colony growth dynamics and phenology.

### Experiment 2 – pollen effects on performances of queenless micro-colonies

To investigate the effects of late-spring pollen on bumble bee performances, we conducted another experiment using queenless microcolonies of *B. impatiens* as *B. griseocollis* colonies did not reach a sufficient size to extract enough workers. These microcolonies were composed only of workers, known to initiate egg laying in the absence of a queen, producing haploid males (Tasei & Aupinel, 2008). We formed 45 micro-colonies of 5 workers placed in plastic containers (15 x 15 x 10 cm, BioBest) at 26 ± 2°C and 55-65% humidity, and randomly attributed to one of three diets. Workers were up to five days old when harvested from the main *B. impatiens* colonies in July (Figure 1; *Start microcolonies*). Microcolonies were fed ad-libitum pollen balls of either 1) commercial wildflower mix (hereafter “CM2”, N=15), 2) commercial mix + hawthorn (*Crataegus canadensis*, hereafter “hawthorn”; N=15), or 3) commercial mix + sumac (*Rhus typhina*, N=15, hereafter “sumac”), as well as 50% sucrose solution. Each microcolony received two pollen balls: a commercial mix pollen ball, as well as a second one of their respective diet treatments (hawthorn or sumac) or another commercial pollen ball (CM2 diet group). Pollen balls were prepared as detailed in experiment 1 and were supplemented to microcolonies every two males produced. Microcolony performance on diet was assessed by recording: (i) number of brood cells and (ii) number of males produced. These traits were recorded every other day for 38 days.

### Pollen source and biochemical characterization

Experimental honey bee collected pollens (red maple, hawthorn and sumac) were obtained frozen from Happy-Culture Inc. (Roxton Falls, QC, Canada) in 2021. Experimental wildflower pollen mixes were obtained from Plant Products (Toronto, ON, Canada). These commercial wildflower mixes are commonly used to rear bumble bees in North-America (Rowe et al., 2023), and thus provide a good control. A multidimensional assessment of the nutrient profile and pollutants of these pollen was conducted as part of previous work (Haroune et al., 2025; Morin et al., 2026). Briefly, this multidimensional assessment consisted in characterizing 32 nutrients in pollen samples by a combination of common assays (Loveridge, Folch, Bradford), HPLC and mass-spectrometry (as previously described (Haroune et al., 2025; Morin et al., 2026)). The botanical origin of all samples was confirmed by microscopic observation (magnification x1000). All samples were maintained on ice during the entire biochemical assessment process to preserve essential nutrients like vitamins. The extended results on pollen composition will not be presented here, as they have been published and are publicly available in an open database (Morin et al., 2026). Data on red maple, hawthorn, sumac and the two commercial mixes will be used in this paper to confront their composition with the performances of *B. impatiens* and *B. griesocollis* founding queens and infer on their nutritional requirements.

### Statistical analyses

#### a) Reproductive outputs of queenright colonies (experiment 1)

We looked at the effects of the diet on *the number of brood cells, workers, males and gynes* produced by founding queens using Generalized Linear Mixed Models (GLMMs) with the *glmmTMB package*, which allows consideration of zero-inflated distribution of data. We applied a truncated negative binomial model for the number of workers (to account for both overdispersion and a marked zero-inflated distribution), a zero-inflated negative binomial for males and brood cells (overdispersion) and a zero-inflated poisson models for gynes (no overdispersion); these distributions best fitted the data in all four cases. We included the diet group and species as fixed effects and the diet*species interactions as an additional fixed effect if its inclusion did not increase the AIC. The site of capture was included as a random effect in all models. We also included queen lifespan as fixed effect for models on the maximum number of brood cells (counted at the end of the experiment) and the number of gynes to account for the fact that some queens had died before reaching colony’s full size or before being able to produce gynes. The significance threshold was set at α<0.05.

#### b) Longevity of founding queens and phenology of colonies (experiment 1)

The longevity of the founding queen (number of days from capture to death) was analyzed using a GLMM (glmmTMB), with a poisson distribution (logit). We included the diet group and the species as fixed effects, and the site of capture as random effect. We also ran GLMMs to look at colonies’ phenology, i.e. the number of workers, males and gynes produced over time. We included the diet group, the species, their interaction and the date as fixed effects. The colony identity and the site of capture were included as random effects.

c) Reproductive outputs of queenless microcolonies (experiment 2)

We modeled the effects of the diet on the maximum number of brood cells and average number of males produced using GLMMs. We used the *glmer* package and fitted a poisson distribution in both cases as there was no overdispersion. We included the diet (hawthorn, mix or sumac), the diet of the colony of origin (red maple or mix, to account for potential carry-over effects) and the diet*initial diet interaction as fixed effects in these two models. We included the colony of origin as random factor to account for repeated measures on workers originating from the same colonies. A Tuckey posthoc test was computed for multiple comparisons.

Analyses were conducted on R Studio (R version 4.4.1), using the *glmmTMB* and *glmer* packages. The significance threshold was set at p < 0.05. Figures were prepared using ggplot2.

## Results

### Experiment 1 – queenright colonies fed red maple and wildflower mix

#### A) Colony strength and colony size

A total of 13 founding queens fed red maple (RM) initiated a colony out of 20 (overall success of initiation of 65%): 8 out of 10 for *B. impatiens* and 5 out of 10 for *B. griseocollis*. In the control group (fed the wildflower commercial mix 1 = CM1), 6 founding queens initiated out of 20 (30% success of initiation): 3 out of 10 for *B. impatiens* and 3 out of 10 for *B. griseocollis*. The *maximum number of brood cells* (colony strength) produced was significantly influenced by queen lifespan (z = 3.35, p=0.001) and the diet*species interaction (z = 2.49, p = 0.013); posthoc analyses highlighted that *B. impatiens* fed RM produced significantly more brood cells than the three other species*diet groups (Figure 2A). Regarding *the total number of workers* produced (colony size), the model revealed a significant effect of the species (z = 5.30, p<0.001, Table S1, Figure S2) but no effect of the diet (z = −1.05, p = 0.29, Figure 2B); the species*diet interaction was excluded from the final model (Table S1).

**Figure 2-.**
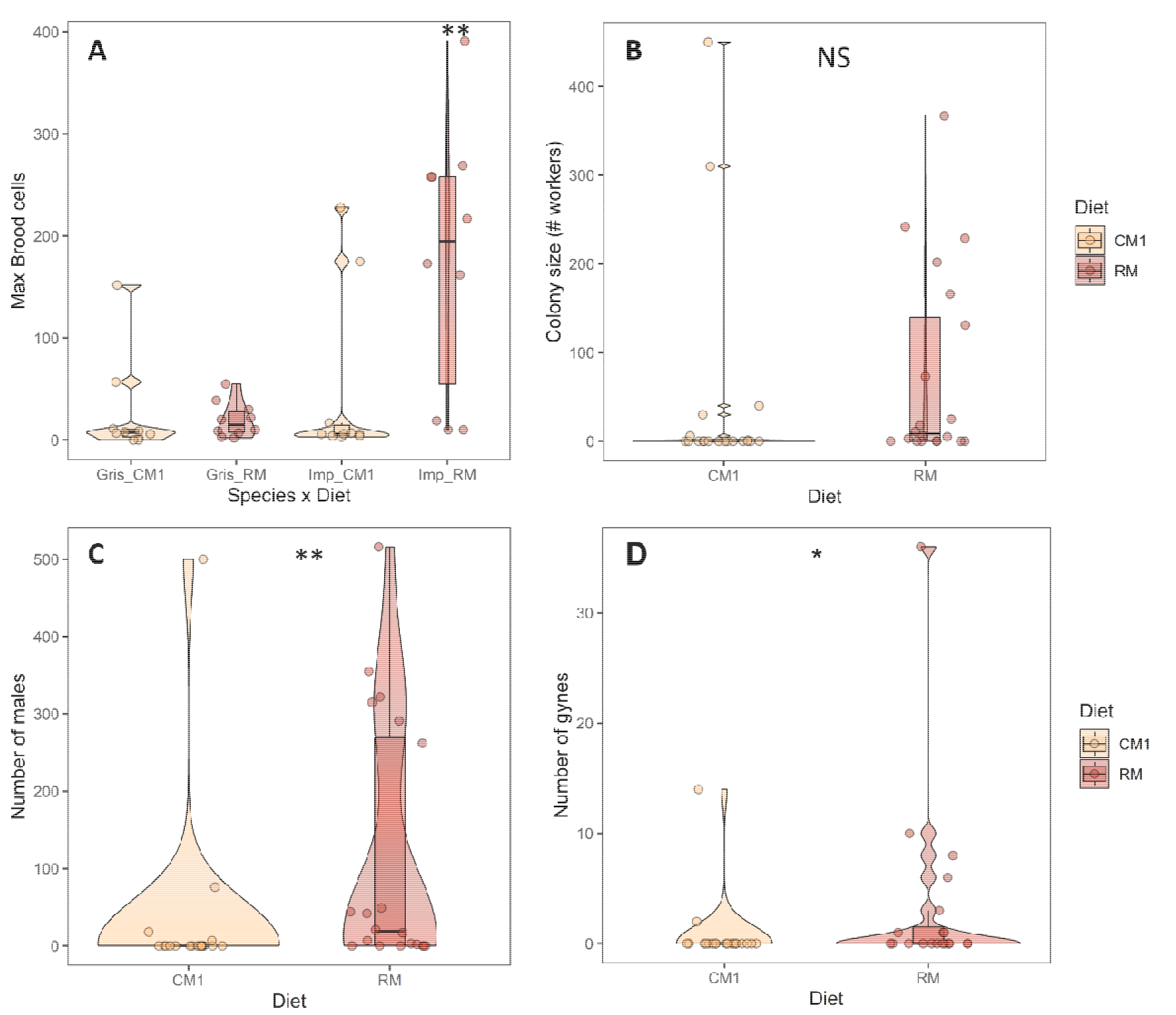
Diet effects on the reproductive output of wild-caught bumble bee queens depending on their diet. In (A) The colony strength (maximum number of brood cells in the colony), depending on the diet and species; RM impatiens queens produced significantly more brood cells than all other groups (p<0.002, **). (B) Colony size (maximum number of workers) depending on the diet group (no diet effect, p>0.05). (C & D) Number of reproductive individuals (males and gynes) produced depending on the diet group (diet effect, males p=0.001, **, gynes p = 0.03, *). Diets: CM1 for commercial mix 1 and RM for red maple 75:25 mix.

#### B) Reproductive individuals

We found an effect of the diet (z = 3.25, p = 0.001) and the species (z = 2.21, p = 0.03) on *the number of males produced*, with more males produced on average in colonies fed RM than in control CM1 colonies (Figure 2C), and by *B. impatiens* than by *B. griseocollis* (emmeans ± SE = 4.42 ± 0.48 and 3.14 ± 0.64, respectively; see Table S2, Figure S3). In total, 689 males were produced in the RM group, compared to 159 males in the control CM1 group.

Similarly, we found a significant effect of the diet (z = 2.14, p = 0.03) and the species (z = 4.31, p <0.001) on *the number of gynes* produced, but no significant effect of queen lifespan (z = 1.77, p = 0.07; Table S3). Overall, more gynes were produced in the RM-fed colonies than in the CM1 ones (Figure 2D). Regarding the species, *B. griseocollis* produced more gynes on average than *B. impatiens* (emmeans ± SE: 0.25±0.99 and −2.37±1.28; Figure S4). In total, 93 gynes were produced in the RM group (N=21 by *B. impatiens* and 72 by *B. griseocollis*), compared to 16 gynes in the CM1 control group (16 by *B. griseocollis*; no *B. impatiens* successfully produced a gyne in this diet group).

#### C) Queen longevity and colonies’ phenology

Founding queens’ lifespan was significantly influenced by the diet (*z-value* = 1.97, p=0.048); on average, queens from the RM group lived longer than queens from the CM1 group (Tuckey Estimate±SE = −0.08±0.04; Table S4). In total, 28 queens (70%) survived until the end of the experiment. The life expectancy of those who did not survive the 130 days, ranged from 16 to 113 days (average 76 days).

GLMMs revealed a significant time*diet interaction on the number of workers produced (*z-value* = 4.41, p<0.001; Table S5): RM queens produced workers earlier in the season than CM1 queens on average (Tuckey, Estimate±SE = −5.15±2.14; Figure 3A; Table S6). The same effect was observed in males (Figure 3B; *z-value* = −3.34, p=0.001) and gynes (Figure 3C; *z-value* = −2.23, p = 0.026; Table S7).

**Figure 3:**
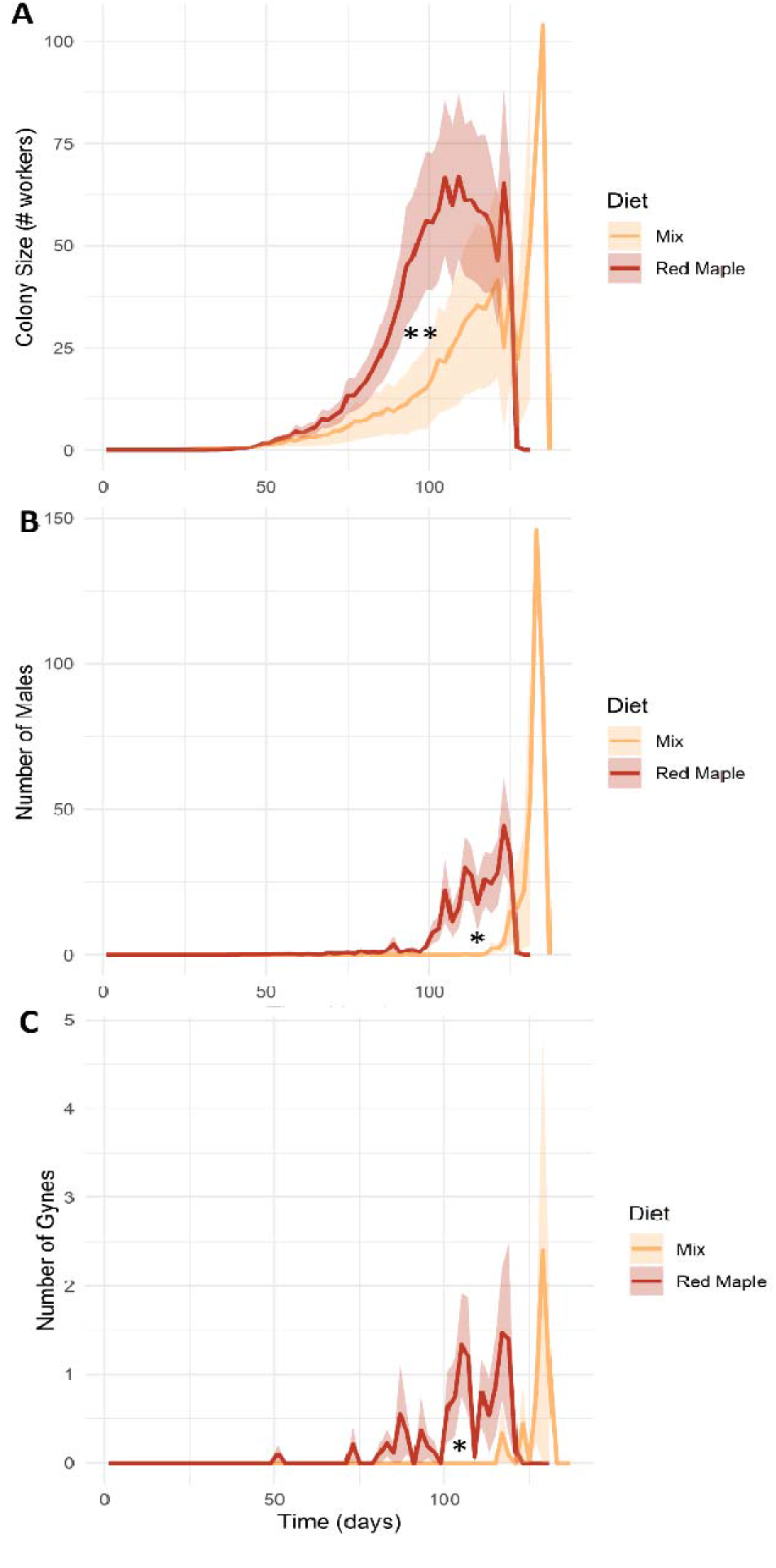
Colony growth over time depending on the diet. In (A) colony size is represented by the number of workers produced over time for each diet group (Mix = CM1, Red Maple = RM). In (B) and (C) the production of reproductive individuals (males and gynes, respectively) by each colony is represented over time as function of diet. We found a significant diet*time interaction (** p<0.01 and * p<0.05). Note that the peak reached at the end for the mix diet group is due to one *B. impatiens* queen that produced a very large number of workers and males (but zero gynes), hence the absence of variation around the line in A and B.

### Experiment 2 – queenless microcolonies fed hawthorn, wildflower mix or sumac pollen

We found a significant effect of the diet on the maximum number of brood cells produced by microcolonies: on average, microcolonies fed with hawthorn performed better than CM2 or sumac microcolonies (z = 2.07, p = 0.04 and z = 2.73, p = 0.006). We also found a strong significant effect of the initial diet (z = 5.35, p <0.001): microcolonies composed of workers from colonies fed RM performed significantly better than workers from colonies fed the CM1 control group in experiment 1 (Figure 4A).

**Figure 4:**
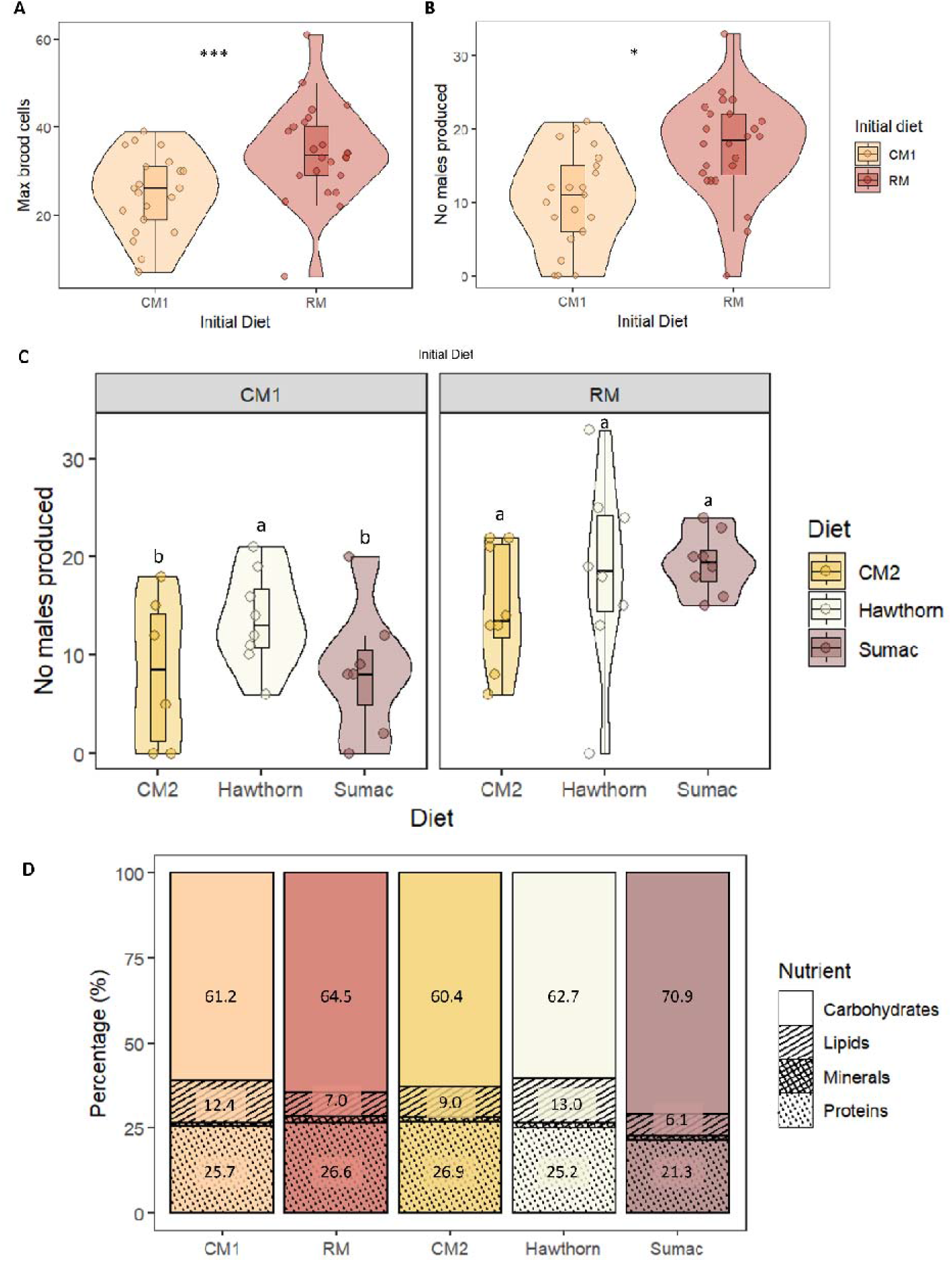
effects of the current and initial diets on the reproductive outputs of queenless microcolonies of Bombus impatiens. The *initial diet* represents the diet provided to the colonies (experiment 1) from which the workers were harvested to form microcolonies (experiment 2), while the *diet* represents the pollen provided to the microcolonies during experiment 2. A) Maximum number of brood cells and B) number of males produced depending on the initial diet. C) number of males produced based on the initial diet (commercial mix 1 = CM1 on the left, Red Maple = RM on the right) and the current respective diets (hawthorn, commercial mix 2 = CM2 and sumac). Different letters and * represent a significant difference at p<0.05; *** for p < 0.001.

The average number of males produced by microcolonies was significantly influenced by the initial diet, the current diet and their interaction (Table S8). Microcolonies formed of workers from RM-fed colonies performed overall better than microcolonies formed of workers from CM1-fed colonies (Figure 4B), with the exception of microcolonies fed hawthorn that performed as good (Figure 4C, post-hoc tests highlighted that CM1-Hawthorn performed better than CM1-CM2 and CM1-Sumac, and as well as RM-Hawthorn, RM-CM2 and RM-Sumac).

### Macro- and micronutrient contents of the diets

Pollen nutrient composition was overall very similar between the five diets, except for vitamins (Figure 4D). Moisture varied between 13.9% (RM pollen) and 14.8% (hawthorn) of fresh pollen mass. Ashes content varied between 2.3% (CM2) and 7.1% (sumac pollen). Regarding carbohydrates, their content varied from 61% (hawthorn pollen) and 71% (sumac pollen). Protein content was lower in sumac pollen (21%) compared to the four other pollens (25-27%). Lipid content varied from 6% (RM pollen) to 13% (hawthorn pollen). Minerals were least abundant in Pollen mix (5.9 ug/g), and most abundant in RM pollen (12.4 ug/g). Regarding vitamins, they were more than twice as abundant in RM and hawthorn than in the three other pollens (Table 1).

**Table 1:**
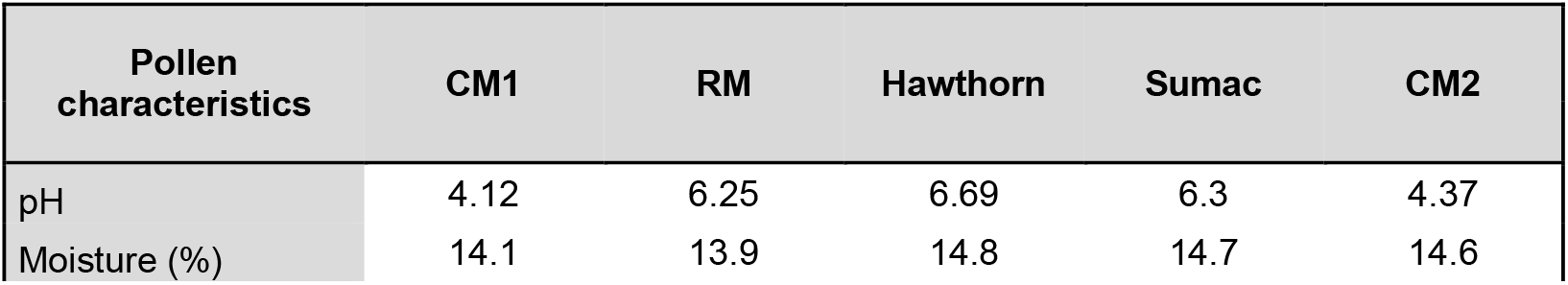
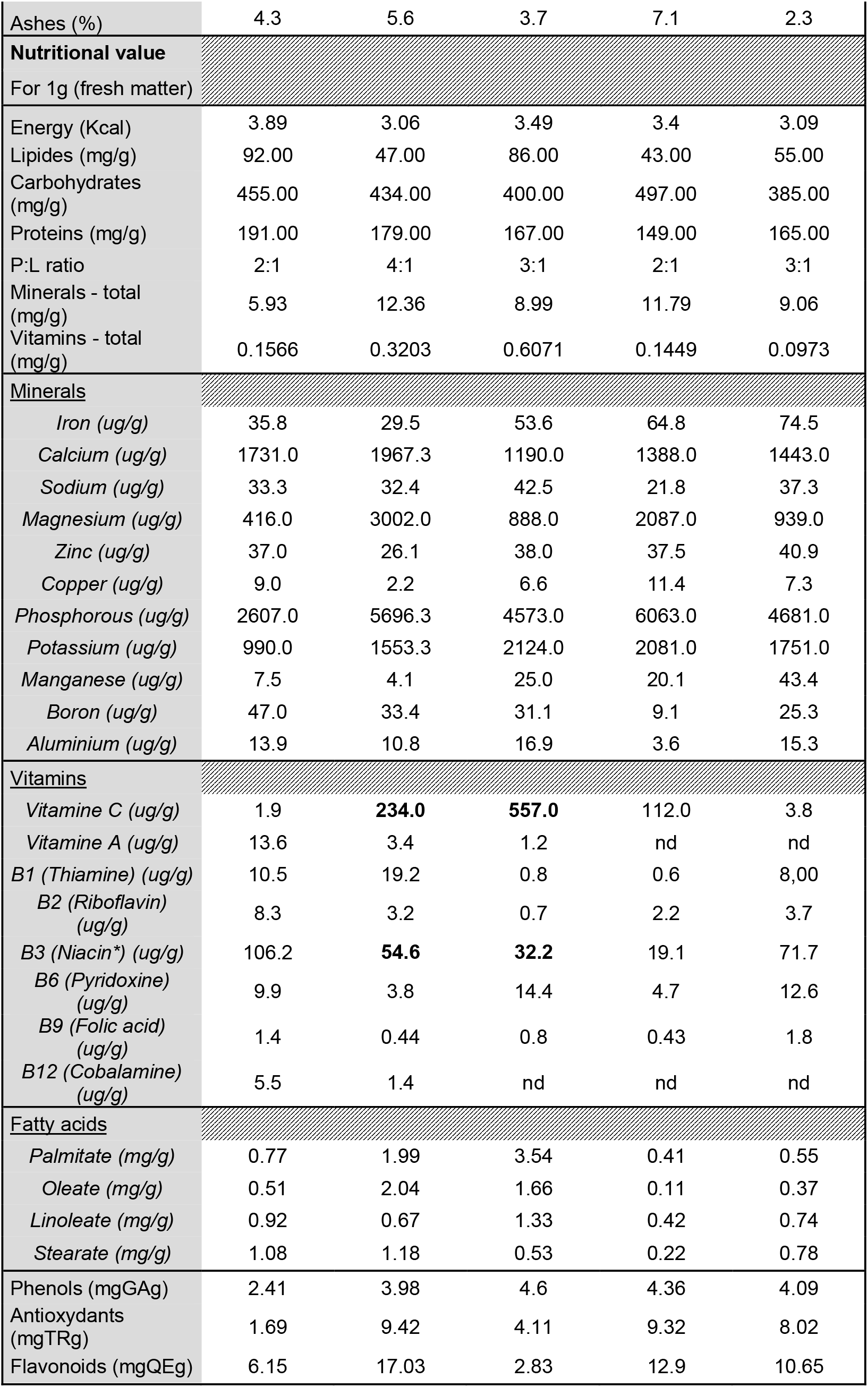
Nutrient composition of the 5 pollen diets provided to bumblebees in experiment 1 and in experiment 2. In experiment 1, CM1 = commercial mix 1, RM = red maple mix; in experiment 2 hawthorn, sumac and commercial mix 2 = CM2. Bold numbers represent nutrients for which we observed a difference of performance in bumble bees.

When focusing on the two diets that significantly improved reproductive performances of founding queens (75:25 Red Maple diet; experiment 1) or queenless microcolonies of workers (hawthorn pollen; experiment 2) we can conclude that a balanced fresh pollen source to support bumble bee reproduction should be composed of 25-27% proteins, 7-13% lipids, 61-64% carbohydrates, 2% minerals, with a moisture ranging from 13.9 to 14.8%. We found no consistent differences in the macronutrient content, protein:lipid ratios (P:L) or minerals between the two diets on which bumble bees performed better (hawthorn and Red Maple diets) and the other pollen, even though RM pollen mix had a the highest P:L ratio (4:1). Hawthorn and RM pollens, however, consistently differed with the three other pollens on two nutrients: vitamin C (ascorbic acid) and vitamin B3 (niacin) as shown in Table 1. Vitamin C was 100 times more abundant in RM pollen than in CM1, and 5 to 145 times more abundant in hawthorn pollen than in sumac and CM2. Vitamin B3 ranged from 32.2 ug/g of pollen in hawthorn and 56.6 ug/g in RM pollen, and was either substantially below (sumac) or above (CM1 and CM2) that range.

## Discussion

Although we initially predicted that pollen from spring-flowering trees would increase reproductive performances of bumble bees compared to a wildflower mix, our results highlighted that this was true for two pollens types – red maple (RM) and hawthorn, which increased queen and workers reproductive success, respectively – while sumac pollen appeared to be limiting the reproduction of queenless microcolonies of workers. While differences in the P:L ratio could explain this difference between sumac and red maple (ratios of 4:1 and 2:1, respectively), it failed to explain the overall differences observed between the five pollens tested. Interestingly, RM pollen had positive carry-over effects on reproductive performances of queenless microcolonies of workers. While looking at the multidimensional nutrient composition of the five pollens tested, vitamin C and vitamin B3 appeared as the two determining factors explaining the benefits of RM and hawthorn pollens on bumble bee reproductive and survival performances.

### Pollen effects on bumble bee performances across experiments

#### Red maple strongly increases queen reproduction

More than twice as many founding queens initiated a colony when fed red maple than fed the commercial wildflower mix (experiment 1). Queens fed RM initiated a colony earlier and produced overall more brood cells but had similar colony sizes than CM1 colonies. Even though the supplementation in RM started in spring and lasted 7±1 weeks (up to July), it had substantial and long-lasting effects on colony performances, including an increased number of reproductive individuals produced in August and September in RM compared to CM1 colonies, an advanced phenology in the production of all castes, increased founding queen longevity, and even carry-over effects on workers reproductive performances (experiment 2; discussed below).

These results have determinant ecological consequences. Indeed, even though we did not find any effect of RM on the average number of workers produced per colony, twice as more workers were produced in *B. impatiens* fed RM pollen than in the CM1 *B. impatiens colonies* in total (1415 over 760). This tendency was not observed in *B. griseocollis*, who produced very small colonies and overall, more gynes than *B. impatiens* (*supplementary information*). Nonetheless, RM-fed colonies still produced four to six times more males and gynes, respectively, than CM1 colonies. Such a difference in the number of adults produced could have key cascading effects on plant-pollinator networks or the provision of pollination services in agroecosystems. Furthermore, previous research in bumble bees showed that low quality food reduced workers’ body size, a trait not recorded in our study, with negative consequences for pollination services (i.e. a reduction in both flower handling efficiency and number of pollen grains deposited on flowers) (Birkenbach et al., 2024). Finally, we also found that RM-fed colonies produced all castes significantly earlier in the season compared to CM1 colonies, which could have crucial consequences on gynes ability to locate a male, copulate, and finding an overwintering site on time before the fall. All these traits are likely to influence bumble bee community persistence over the years in the wild as well as pollination services in agroecosystems.

#### Hawthorn improve success of microcolonies with carryover effects from red maple pollen

Experiment 2 on queenless microcolonies of workers revealed an interesting interaction between their current diet (pollen diets provided to workers, i.e. hawthorn, sumac or commercial mix 2; CM2) and their initial diet (pollen provided to founding queens, i.e., red maple and commercial mix 1; CM1). Overall, workers’ microcolonies with RM as initial diet performed better than microcolonies harvested from CM1 (control) colonies. When looking only at microcolonies with CM1 as initial diet, the ones provided with hawthorn as current pollen diet performed better than the ones provided with sumac and CM2, and as well as all the microcolonies with RM as initial diet. This highlights that the current diet interacted with the initial diet, and that only hawthorn allowed microcolonies to “recover” from the initial CM1 diet which did not meet nutritional requirements of colonies (as discussed above). This also highlights positive carryover effects of RM pollen on workers, i.e. influencing reproductive outputs and population dynamics through time. In this context, it not only reveals that RM pollen increased reproductive abilities of founding queens but also improved the reproductive capacities of their offspring – here workers – even after they were switched to alternative pollen diets. So far, the rare studies considering ecological carryover effects in bees highlighted negative effects of pollen shortage (e.g. in honey bees; (Requier et al., 2017)) or of pesticides (e.g. in solitary bees; (Stuligross & Williams, 2021)). Here, however, we highlight positive carryover effects of RM pollen. Considering that diets can offset negative effects of pesticides or heatwaves on bees under certain circumstances (Vanderplanck et al., 2019), future work should investigate the ability of some nutritious pollen – like red maple, hawthorn or pollen with similar nutritional signature (Morin et al., 2026) – to offset such negative carryover effects. In our study, however, carryover effects were recorded on queenless microcolonies of workers – which is not the reproductive caste in a natural setting. However, workers are known to influence the physiology and fecundity of the founding queens (Sarro et al., 2021). Moreover, if the founding queen dies prematurely – i.e. before the emergence of reproductives – workers will take over, care for the brood and produce males (Goulson, 2010), ensuring the persistence of the colony and emergence of reproductives. Our results suggest that workers may better perform at this task if fed with nutritious red maple and hawthorn pollen, respectively flowering in early spring and early summer. A diet consisting of these two pollens, which bloom successively in the spring and early summer, could provide a good nutritional balance that supports the seasonal needs of bumblebees.

Importantly, even though using queenless microcolonies or workers is a commonly used method to estimate whole-colony reproduction as a function of diet (Giacomini et al., 2018; Moerman et al., 2016, 2017), it may not fully mimic responses of queens and gynes, potentially underestimating the negative effects of dietary deficiencies (Wynants et al., 2022). Despite strong similarities between nutrient content in CM1 and CM2 pollens, CM1 strongly limited reproductive success of founding queens in experiment 1, since most queens did not produce a colony or reproductive individuals; in experiment 2, although we found a diet effect, all microcolonies performed relatively well in producing eggs and males, even on the less nutritious CM2. Based on these results and since queenright colonies are more sensitive to nutrient deficiencies than microcolonies of workers (Wynants et al., 2022), we recommend focusing on queenright colonies to assess the nutritional requirements of bumble bees in the future. To fully capture how pollen harvested by founding queens and workers will affect next generations, future experiments should thus consider looking at carryover effects of pollen provided to queenright colonies on the fitness of reproductive individuals, by measuring how pollen provided to founding queens and to colonies affect life-history traits (adult body size/mass, overwintering survival, reproductive phenology and success the next year for gynes, and traits such as body mass/size, longevity or sperm motility in males) or nutrition-related traits (e.g. protein or glycogen contents) of reproductive casts (Woodard et al., 2019).

#### Ecological implications of hawthorn and red maple pollen for wild bumble bees

The benefits of red maple pollen on queen and colony performances, or of hawthorn pollen on workers’, could be subjected to external factors not considered in this study, conducted under controlled conditions and focusing on basal nutritional requirements. For instance, queen reproductive outputs are known to be influenced by El Nino years in South-America – and more broadly by weather (Riaño-Jiménez et al., 2020), while the quality of red maple pollen varies between years, potentially due to masting cycles (Morin et al., 2026). Furthermore, recent work highlighted that low food quality alone did not influence the number of workers produced by European *B. terrestris* colonies – which is consistent with our findings – while when interacting with pesticides, it negatively influenced the number of workers produced (Birkenbach et al., 2024). It is thus likely that the differences that we recorded between RM- and CM-fed colonies would be exacerbated in the wild, where colonies are exposed to cocktails of pesticides and other Anthropogenic stressors. This further supports the need to ensure access to high-quality food for wild pollinators in agroecosystems ((Birkenbach et al., 2024; Vaudo et al., 2018), reviewed in (Tissier et al., 2025)). Importantly, the nutritional needs of bumble bees might vary depending on their physiological state and health status. For instance, nutritional-rich pollen like red maple may support reproduction and colony development, but can also favor the development of some gut parasites like *Crithidia bombi*, a trypanosome infecting bumble bees (Figueroa et al., 2023). On the contrary, Asteraceae pollen – like sunflower, dandelion or goldenrod – have antimicrobial properties that can be beneficial to infected bees (Figueroa et al., 2023; Genissel et al., 2002; Giacomini et al., 2018; LoCascio et al., 2019). Such Asteraceae pollens, however, generally do not support bumble bee reproduction because of a lack of proteins (≤16%), reduce longevity and may cause abnormal behaviors owing to a deficiency in tryptophan and vitamin B3 (Genissel et al., 2002; McAulay & Forrest, 2019; Tissier et al., 2023). There might thus be a trade-off– i.e. the need for a balance between nutritional and antimicrobial pollens– depending on the life-cycle stage and the infection status. More studies are thus needed to better assess the breadth of nutritional needs supporting bumble bee queen performance and health in changing environments where they are exposed to multiple stressors.

### Effects of vitamins on bumble bee performances and expected ecological consequences

#### Vitamins B3 and C best explain observed differences in reproductive success than P:L ratios

Previous research revealed that the best P:L ratios supporting bee reproductive and survival performances were between 4:1 and 5:1 (Vaudo et al., 2016, 2018) or that bees need around 20% proteins in their diet (Barraud et al., 2022). While sumac pollen was the lowest in proteins (21% of fresh mass), protein content of commercial pollen mixes made up more than 25% of the fresh mass. When looking at P:L ratios, however, both CM1 and sumac were less performant (with ratios of 2:1) than hawthorn, RM and CM2 – which could partially explain the lower performances of colonies and microcolonies on these diets. Nonetheless, when comparing CM2 (low performance of microcolonies) and hawthorn, both having P:L ratio is of 3:1, we fail to explain the differences observed in their respective reproductive performances. Overall, when comparing all diets, P:L ratios alone are not sufficient to explain differences between the commercial pollen mixes and the two diets on which bumble bees performed best (hawthorn and RM). This highlights that when the diet contains sufficient proteins, other nutrients may become limiting, such as vitamins. Our results showed that vitamin C and vitamin B3 appeared as the two common factors differentiating pollens on which bumble bees performed well (hawthorn and RM) from sumac and commercial pollen mixes.

Vitamin C supplementation has a protective effect on oxidative stress, increasing performances of *A. mellifera carnica* (Farjan et al., 2012). In this study, oxidative stress was found to be higher after workers emergence, and vitamin C played a protective role at that life-stage, leading to a 33% reduction of bee losses over winter. Positive effects on brood production were recorded when diet was supplemented by 500 ug/g of vitamin C (Herbert et al., 1985), which is equivalent to what was provided by red maple and hawthorn pollen (which varied between 234-557 ug/g). Those early studies on the benefits of pollen vitamin C on brood rearing in honey bees also reveal a seasonal trend, with pollen collected in spring (May) having significantly greater content in vitamin C than pollens collected in summer (Aug). This support the idea that early-flowering pollens are of key importance for wild bee reproduction – by providing key nutrients for brood development that later become scarce in the environment, like vitamins.

Vitamin B3 is essential in all animals and is known to be limiting reproduction in vertebrates even when dietary proteins are abundant (Tissier et al., 2017, 2023). Previous studies highlighted that honey bees actively regulate their vitamin B3 intake, or that of its precursor, the essential amino acid tryptophan, with direct consequences both on macronutrient and daily food intake as well as survival (Elsayeh et al., 2022; Fengkui et al., 2015). Moreover, although the nutrient content of pollen collected by honey bee worker bees greatly varies (from 30-210 ug/g of pollen), niacin content in royal jelly –provided to larvae of future queens– is strongly regulated and varies only between 42-88 ug/g (Elsayeh et al., 2022). This is closer to the range of 32.2 ug/g in hawthorn and 54.6 ug/g in red maple that we report in this study, with the three other pollens –on which bumble bees did not perform well– exhibiting niacin content of 19.1 ug/g (substantially lower) or of 71.7 ug/g and 106.2 ug/g (substantially greater). Our results thus suggest that vitamin B3 intake, as shown for honey bees, is tightly regulated in bumble bees and that an appropriated range of 32.2-54.6 ug/g is adapted for the reproductive performances of *B. impatiens* (Experiments 1 and 2), while for *B. griseocollis* a content of 54.6 ug/g seem adapted to support queen reproduction and colony growth (Experiment 1). Determining the appropriate vitamin B3 intake of various bumble bee species could be particularly important for proper development of gynes, as is the 42-88 ug/g observed in royal jelly of honey bees (Elsayeh et al., 2022).

#### Beyond vitamins and towards interactive effects with pollutants in natural settings

It is worth noting that beyond the 32 nutrients that were assessed in this study, other factors could play a role on success, such as content in essential amino acids (EAAs). Total EAA content of the diet is known to be a determining factor for bumble bee reproductive performances (Barraud et al., 2022; Moerman et al., 2016, 2017). Maple pollens are known to have an elevated total content in EAAs and to be particularly rich in the precursor of vitamin B3, the EAA tryptophan (Chrzanowska et al., 2024). Assessing the proper EAA requirements of various bumble bee species could thus be the focus of future studies.

While pollen pollutants could also be proposed as a potential explanation for pollen effects on reproductive performances, previous work investigating 100 pollutants (pesticides, pharmaceuticals and heavy metals) highlighted reduced contamination levels of the pollen used in this study (Morin et al., 2026). Briefly, red maple had a greater content in heavy metals (especially Arsenic, at 3.39 ug/g and Cadmium at 1.99 ug/g) compared to other pollens. Low levels of atrazine were found in sumac and hawthorn pollens (<0.25 ng/g), and we found ≤32.1 ng/g of glyphosate in sumac and CM1 pollens (provided to both CM and RM colonies for most of the study). Apart from these, all metabolites of pesticides and pharmaceuticals were below the limit of detection (Haroune et al., 2025; Morin et al., 2026). It is thus unlikely that pollutants played a key role in explaining the differences in reproductive performances between pollens in this study. The main concerns would come from glyphosate, though the levels to which bumble bees were exposed in this study through CM1 pollen (32.1 ng/g) were much lower than the 5-39 ug/g residues found in pollen in other studies (Thompson et al., 2022) and those shown to have sublethal effects on bees (in the order of ug/g of food; (Klaus et al., 2021; Weidenmüller et al., 2022). Even though neonicotinoid insecticides are known to affect reproductive performances of solitary bees (*Osmia bicornis*) at 3.00 ng/g (Klaus et al., 2021), it is unclear whether glyphosate may affect the reproductive performances of bumble bees at the concentration found in our pollen (ng/g), since studies investigating the sublethal effects of this herbicide have used greater concentrations (ug/g). Furthermore, although RM-fed queens were provided red maple pollen for 7±1 weeks, they were mostly fed on CM1 throughout the 130-days experiment. Indeed, RM-fed queens received a single pollen ball of 75% red maple and 25% commercial mix during the experimental phase of 7±1 weeks, then all additional pollen balls provided after worker emergence to ensure unlimited access to pollen (from July to September) were composed of CM1, as was the pollen provided during installation. Queens and colonies fed RM were thus also exposed to glyphosate through CM1 pollen throughout the experiment. We cannot rule-out, however, that the nutritional value of red maple pollen somehow offset potential sublethal effects of glyphosate. Previous studies indeed showed that vitamins, particularly vitamin C and B-complex vitamins, have protective effects against pollutants, in bees exposed to pesticides (EL-Gendy et al., 2010) and mammals exposed to heavy metals like cadmium (Abdelaziz et al., 2013). Taken altogether, this confirm the benefits of red maple and hawthorn for bumble bee reproductive performance and longevity, despite the potential exposure to some pollutants, which is likely to happen in a natural context (Knauer et al., 2025).

## Conclusion

Our results bring new knowledge improving the global understanding of bumble bee nutritional needs, especially the requirements of wild overwintered founding queens. These results will benefit conservation breeding programs by providing detailed information on nutritional needs to successfully breed bumble bees in conservation breeding programs, while providing useful nutritional information to design conservation habitats supporting wild bee populations. Since bee nutritional requirements may be subjected to various external factors – such as infections, pesticides or heat waves – future studies should investigate the nutritional needs of bumble bee queens submitted to such environmental stressors to better assess the breadth of nutritional needs supporting their performance and health and inform conservation management strategies.

## Acknowledgements

We than the Liber Ero foundation and fellows for their support. We thank Yann Loranger and Isabelle Rabbat from Happy-Culture inc. for providing red maple, sumac and hawthorn pollen used in this study. Many thanks to Ellen Richard from WPC for her help with the experiment on microcolonies, and to Genevieve Rowe for her advice. We dedicate this article to our forever remembered mentor, Sheila R. Colla.

