## Supplementary material for "Vitamins in native tree pollens are essential to support fitness of wild-caught *Bombus* spp. queens, with positive carry-over effects": Table S1

Mathilde L. Tissier, and Sarah MacKell
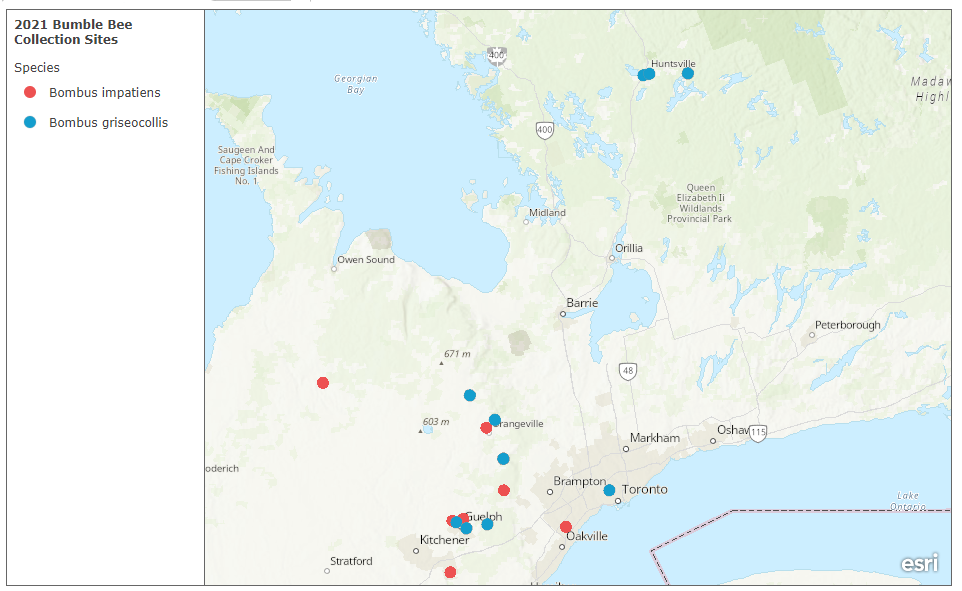

**Figure S1:** Map of the 19 sites (Sounthern Ontario, Canada) where bumble bee queens were collected for the experiment.

**Table S1:** output of the model looking at the effect of the diet on the number of workers produced in each diet group (experiment 1). Model structure: Workers = glmmTMB(TotalNoWork ~ Species + DietGroup + (1|Site_area), data=Data_final, zi =~1, family = truncated_nbinom2()).

|  | **Total Number of workers** | | | | |
| --- | --- | --- | --- | --- | --- |
| *Predictors* | *Estimate* | *std. Error* | *CI* | *Z-value* | *p* |
| **Count Model** | | | | | |
| (Intercept) | 3.07 | 0.47 | 2.14 – 4.00 | 6.50 | **<0.001** |
| Species [Impatiens] | 2.59 | 0.49 | 1.63 – 3.55 | 5.30 | **<0.001** |
| DietGroup [RM] | -0.54 | 0.51 | -1.55 – 0.47 | -1.05 | 0.295 |
| (Intercept) | 0.98 |  | 0.48 – 2.02 |  |  |
| **Zero-Inflated Model** | | | | | |
| (Intercept) | 0.10 | 0.32 | -0.52 – 0.72 | 0.32 | 0.752 |
| **Random Effects** | | | | | |
| σ^2^ | 0.71 | | | | |
| τ_00_ _Site_area_ | 0.00 | | | | |
| N _Site_area_ | 9 | | | | |
| Observations | 40 | | | | |
| Marginal R^2^ / Conditional R^2^ | 0.718 / NA | | | | |

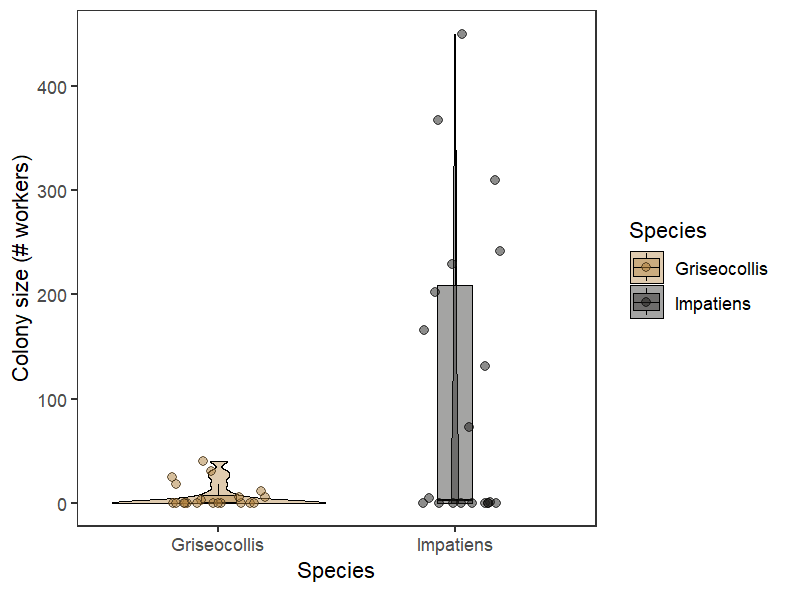

**Figure S2: Colony size depending on the species.** The total number of workers produced over the season is shown for *B. griseocollis* and *B. impatiens* (species effect, p<0.05; see Table S1 above).

| **Table S2**: output of the model looking at the effect of the diet on the number of males produced in each diet group (experiment 1). Model structure: Males = glmmTMB (TotalNoMales ~ Species + DietGroup + (1\|Site_area), data=Data_final, zi =~1, family = nbinom1()). | | | | | |
| --- | --- | --- | --- | --- | --- |
|  | **Total Number of Males** | | | | |
| *Predictors* | *Estimate* | *std. Error* | *CI* | *Statistic* | *p* |
| **Count Model** | | | | | |
| (Intercept) | 2.05 | 0.83 | 0.42 – 3.68 | 2.46 | **0.014** |
| Species [Impatiens] | 1.28 | 0.58 | 0.14 – 2.42 | 2.21 | **0.027** |
| DietGroup [RM] | 2.18 | 0.67 | 0.87 – 3.49 | 3.25 | **0.001** |
| (Intercept) | 234.72 |  | 83.01 – 663.69 |  |  |
| **Zero-Inflated Model** | | | | | |
| (Intercept) | -1.42 | 0.82 | -3.02 – 0.18 | -1.74 | 0.082 |
| **Random Effects** | | | | | |
| σ^2^ | 0.01 | | | | |
| τ_00_ _Site_area_ | 0.41 | | | | |
| ICC | 0.98 | | | | |
| N _Site_area_ | 9 | | | | |
| Observations | 40 | | | | |
| Marginal R^2^ / Conditional R^2^ | 0.797 / 0.997 | | | | |

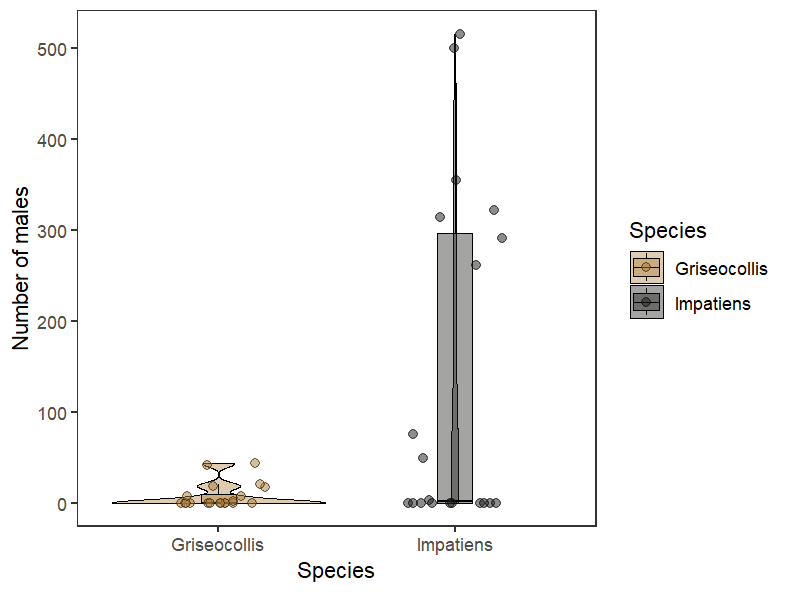
**Figure S3: Number of males produced depending on the species.** The total number of males produced by the end of the experiment is shown for *B. griseocollis* and *B. impatiens* (species effect, p<0.05; see table S2 above).

| **Table S3**: output of the model looking at the effect of the diet on the number of gynes produced in each diet group (experiment 1). Model structure: Gynes = glmmTMB (glmmTMB(TotalNoGynes ~ Species + DietGroup + QueenLifespan + (1\|Site_area), data=Data_final, zi =~1, family = poisson())). | | | | | |
| --- | --- | --- | --- | --- | --- |
|  | **TotalNoGynes** | | | | |
| *Predictors* | *Estimate* | *std. Error* | *CI* | *Statistic* | *p* |
| **Count Model** | | | | | |
| (Intercept) | -7.87 | 4.37 | -16.43 – 0.69 | -1.80 | 0.071 |
| Species [Impatiens] | -2.61 | 0.61 | -3.80 – -1.42 | -4.31 | **<0.001** |
| DietGroup [RM] | 3.32 | 1.55 | 0.28 – 6.37 | 2.14 | **0.033** |
| QueenLifespan | 0.06 | 0.03 | -0.01 – 0.12 | 1.77 | 0.077 |
| **Zero-Inflated Model** | | | | | |
| (Intercept) | -0.60 | 0.63 | -1.83 – 0.62 | -0.97 | 0.333 |
| **Random Effects** | | | | | |
| σ^2^ | 0.59 | | | | |
| τ_00_ _Site_area_ | 3.64 | | | | |
| ICC | 0.86 | | | | |
| N _Site_area_ | 9 | | | | |
| Observations | 40 | | | | |
| Marginal R^2^ / Conditional R^2^ | 0.623 / 0.947 | | | | |

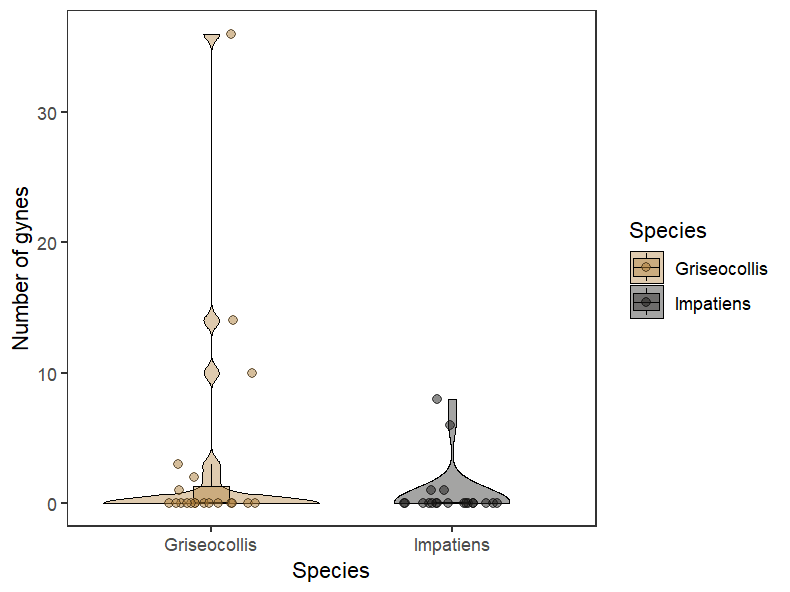

**Figure S4: Number of gynes produced depending on the species.** The total number of gynes produced by the end of the experiment is shown for *B. griseocollis* and *B. impatiens* (species effect, p<0.05; see table S3 above).

**Table S4**: output of the model looking at the effect of the diet on the longevity of the founding queen (experiment 1). Model structure: glmmTMB(QueenLifespan ~ Species + DietGroup + (1|Site_area), data=Data_final, family = poisson()). Since *B. griseocollis* queens were installed later than *B. impatiens* queens owing to their respective phenology in the wild (see the main text) the species effect is not discussed in the manuscript as it may be an artefact of their difference in emergence dates.

|  | **QueenLifespan** | | | | |
| --- | --- | --- | --- | --- | --- |
| *Predictors* | *Estimate* | *std. Error* | *CI* | *Z-value* | *p* |
| (Intercept) | 4.56 | 0.06 | 4.44 – 4.68 | 76.08 | **<0.001** |
| Species [Impatiens] | 0.19 | 0.03 | 0.12 – 0.25 | 5.51 | **<0.001** |
| DietGroup [RM] | 0.08 | 0.04 | 0.02 – 0.15 | 1.97 | **0.048** |
| **Random Effects** | | | | | |
| σ^2^ | 0.01 | | | | |
| τ_00_ _Site_area_ | 0.02 | | | | |
| ICC | 0.73 | | | | |
| N _Site_area_ | 9 | | | | |
| Observations | 40 | | | | |
| Marginal R^2^ / Conditional R^2^ | 0.243 / 0.793 | | | | |

**Table S5**: output of the model looking at the effect of the diet on the phenology of worker production in the colonies (experiment 1). Model structure: PhenoWork1 = glmmTMB(No_workers ~ DietGroup + Day + DietGroup:Day + Species + (1|Bee_ID) + (1|Site_area), data=Temporal_Data_2021, zi =~1, family = nbinom2()).

|  | **Number of workers** | | | | |
| --- | --- | --- | --- | --- | --- |
| *Predictors* | *Estimate* | *std. Error* | *CI* | *Z-value* | *p* |
| **Count Model** | | | | | |
| (Intercept) | -11.84 | 2.22 | -16.19 – -7.48 | -5.33 | **<0.001** |
| Diet Group [RM] | 4.63 | 2.14 | 0.43 – 8.83 | 2.16 | **0.031** |
| Day | 0.05 | 0.00 | 0.05 – 0.05 | 31.36 | **<0.001** |
| Species [Impatiens] | 1.87 | 2.01 | -2.07 – 5.81 | 0.93 | 0.352 |
| Diet Group [RM] × Day | 0.01 | 0.00 | 0.01 – 0.01 | 4.41 | **<0.001** |
| (Intercept) | 3.91 |  | 3.28 – 4.65 |  |  |
| **Zero-Inflated Model** | | | | | |
| (Intercept) | -2.83 | 0.27 | -3.35 – -2.31 | -10.65 | **<0.001** |
| **Random Effects** | | | | | |
| σ^2^ | 0.00 | | | | |
| τ_00_ _Bee_ID_ | 31.86 | | | | |
| τ_00_ _Site_area_ | 0.00 | | | | |
| ICC | 1.00 | | | | |
| N _Bee_ID_ | 41 | | | | |
| N _Site_area_ | 9 | | | | |
| Observations | 2192 | | | | |
| Marginal R^2^ / Conditional R^2^ | 0.274 / 1.000 | | | | |

**Table S6**: output of the model looking at the effect of the diet on the phenology of male production in the colonies (experiment 1). Model structure: glmmTMB(Tot_No_Males ~ Diet_Group + Day + Diet_Group:Day + Species + (1|Bee_ID) + (1|Site_area), data=Temporal_Data_2021, zi =~1, family = nbinom2()).

|  | **Number of males** | | | | |
| --- | --- | --- | --- | --- | --- |
| *Predictors* | *Estimate* | *std. Error* | *CI* | *Z-value* | *p* |
| **Count Model** | | | | | |
| (Intercept) | -24.00 | 3.38 | -30.62 – -17.38 | -7.11 | **<0.001** |
| Diet Group [RM] | 15.61 | 3.51 | 8.72 – 22.49 | 4.44 | **<0.001** |
| Day | 0.18 | 0.03 | 0.13 – 0.23 | 7.01 | **<0.001** |
| Species [Impatiens] | -0.30 | 1.25 | -2.76 – 2.15 | -0.24 | 0.810 |
| Diet Group [RM] × Day | -0.09 | 0.03 | -0.14 – -0.04 | -3.34 | **0.001** |
| (Intercept) | 0.53 |  | 0.29 – 0.95 |  |  |
| **Zero-Inflated Model** | | | | | |
| (Intercept) | -0.80 | 0.41 | -1.61 – -0.00 | -1.97 | **0.049** |
| **Random Effects** | | | | | |
| σ^2^ | 0.00 | | | | |
| τ_00_ _Bee_ID_ | 7.78 | | | | |
| τ_00_ _Site_area_ | 1.80 | | | | |
| ICC | 1.00 | | | | |
| N _Bee_ID_ | 41 | | | | |
| N _Site_area_ | 9 | | | | |
| Observations | 2208 | | | | |
| Marginal R^2^ / Conditional R^2^ | 0.849 / 1.000 | | | | |

**Table S7**: output of the model looking at the effect of the diet on the phenology of gyne production in the colonies (experiment 1). Model structure: glmmTMB(Tot_No_Gynes ~ Diet_Group + Day + Diet_Group:Day + Species + (1|Bee_ID) + (1|Site_area), data=Temporal_Data_2021, zi =~1, family = nbinom1()).

|  | **Number of gynes** | | | | |
| --- | --- | --- | --- | --- | --- |
| *Predictors* | *Estimate* | *std. Error* | *CI* | *Z value* | *p* |
| **Count Model** | | | | | |
| (Intercept) | -24.73 | 6.77 | -38.01 – -11.46 | -3.65 | **<0.001** |
| Diet Group [RM] | 17.21 | 6.77 | 3.95 – 30.47 | 2.54 | **0.011** |
| Day | 0.18 | 0.05 | 0.08 – 0.28 | 3.37 | **0.001** |
| Species [Impatiens] | -2.49 | 1.01 | -4.47 – -0.52 | -2.47 | **0.013** |
| Diet Group [RM] × Day | -0.12 | 0.05 | -0.23 – -0.01 | -2.23 | **0.026** |
| (Intercept) | 4.02 |  | 2.35 – 6.87 |  |  |
| **Zero-Inflated Model** | | | | | |
| (Intercept) | -17.88 | 6260.17 | -12287.58 – 12251.82 | -0.00 | 0.998 |
| **Random Effects** | | | | | |
| σ^2^ | 1.14 | | | | |
| τ_00_ _Bee_ID_ | 3.04 | | | | |
| τ_00_ _Site_area_ | 0.58 | | | | |
| ICC | 0.76 | | | | |
| N _Bee_ID_ | 41 | | | | |
| N _Site_area_ | 9 | | | | |
| Observations | 2208 | | | | |
| Marginal R^2^ / Conditional R^2^ | 0.914 / 0.979 | | | | |

**Table S8**: output of the model looking at the effect of the initial diet, the diet and their interaction on the production of males by queenless microcolonies of workers (experiment 2). Model structure: glmer(No_males_tot ~ Diet + Initial_Diet + Diet*Initial_Diet + (1|Initial_Colony), data=Microcolony_Data, family = poisson()).

|  | **Number of males** | | | | |
| --- | --- | --- | --- | --- | --- |
| *Predictors* | *Estimate* | *std. Error* | *CI* | *Statistic* | *p* |
| (Intercept) | 2.61 | 0.10 | 2.42 – 2.80 | 27.27 | **<0.001** |
| Diet [CM2] | -0.49 | 0.17 | -0.83 – -0.16 | -2.88 | **0.004** |
| Diet [Sumac] | -0.48 | 0.16 | -0.80 – -0.16 | -2.97 | **0.003** |
| Initial Diet [RM] | 0.30 | 0.13 | 0.05 – 0.55 | 2.37 | **0.018** |
| Diet [CM2] × Initial Diet [RM] | 0.28 | 0.21 | -0.13 – 0.69 | 1.33 | 0.183 |
| Diet [Sumac] × Initial Diet [RM] | 0.53 | 0.20 | 0.14 – 0.92 | 2.69 | **0.007** |
